# Endogenous miRNA sponges mediate the generation of oscillatory dynamics for a non-coding RNA network

**DOI:** 10.1101/292029

**Authors:** Andrew Dhawan, Adrian L. Harris, Francesca M. Buffa, Jacob G. Scott

**Affiliations:** Department of Oncology, University of Oxford, Oxford, United Kingdom; Departments of Translational Hematology and Oncology Research, Cleveland Clinic, Cleveland, United States

## Abstract

Oscillations are crucial to the sustenance of living organisms, across a wide variety of biological processes. In eukaryotes, oscillatory dynamics are thought to arise from interactions at the protein and RNA levels; however, the role of non-coding RNA in regulating these dynamics remains understudied. In this work, using a mathematical model, we show how non-coding RNA acting as microRNA (miRNA) sponges in a conserved miRNA - transcription factor feedback motif, can give rise to oscillatory behaviour. Control of these non-coding RNA can dynamically create oscillations or stability, and we show how this behaviour predisposes to oscillations in the stochastic limit. These results, supported by emerging evidence for the role of miRNA sponges in development, point towards key roles of different species of miRNA sponges, such as circular RNA, potentially in the maintenance of yet unexplained oscillatory behaviour. These results help to provide a paradigm for understanding functional differences between the many redundant, but distinct RNA species thought to act as miRNA sponges in nature, such as long non-coding RNA, pseudogenes, competing mRNA, circular RNA, and 3’ UTRs.

**Author summary:** We analyze the effects of a newly discovered species of non-coding RNA, acting as microRNA (miRNA) sponges, on intracellular signalling dynamics. We show that oscillatory behaviour can arise in a time-varying manner in an over-represented transcriptional feedback network. These results point towards novel hypotheses for the roles of different species of miRNA sponges, such as their increasingly understood role in neural development.

## Introduction

Oscillations are intrinsic to the behaviour of biological systems, across scales, species, stages of development, and in health and disease [1–3]. For example, during organismal development, oscillations are crucial to the generation of vertebrae, in a process termed somitogenesis [4-8]. During this stage of development, embryonic cells entrain synchronised oscillations, resulting in the development of vertebrae in a coordinated, clock-like process. In organisms exhibiting circadian rhythms, synchronised patterns of neurotransmitter and neurohormonal release are coupled to oscillatory modes [9–11]. For both of these cases, a fundamental question is how a complex interacting system of biomolecules, with intrinsic stochasticity and uncertainty, is able to produce and sustain oscillatory behaviour. In somitogenesis, a seminal work in mathematical biology has proposed the ‘clock and wavefront’ model, which predicts the occurrence of oscillations arising from a biochemical network and diffusive effects [12–15]. Likewise, the Hes1 transcription factor involved in stem cell differentiation participates in a negative feedback loop and has been shown to exhibit stochastic, diffusion-driven oscillatory behaviour [16–18]. For circadian oscillators, the discovery of the regulation of the Period protein and intercellular coupling has shown how oscillations can emerge [3, 10, 11]. Thus, oscillatory behaviour arises in these systems from carefully balanced interactions at the RNA and protein level, among species with specific kinetic properties, giving rise to tunable, dynamic oscillations, even in a noisy biological environment.

Recently, the catalogue of RNA species participating in the dynamics of the transcriptome has been expanded significantly, with the discovery of non-coding RNAs (ncRNAs) - RNA that do not appear to be protein-coding. The manner in which the various species of ncRNAs affect these oscillatory dynamics, if they do at all, is to be determined, as predicted functions remain elusive for circular RNA (circRNA), long non-coding RNA (lncRNA), and pseudogenes [19–22]. One common trait among each of these ncRNA is thought to be the competitive binding of microRNA (miRNA), inhibiting the ability of miRNA to bind mRNA [23]. This competition for miRNA binding is termed ‘sponging’, and is thought to be a primary function of certain circRNA, pseudogenes, expressed 3’ UTRs, and potentially a function for lncRNA as well, as identified through sequence complementarity [20]. In this work, we model these ncRNA, acting as a generalised miRNA sponge on an over-represented miRNA-mRNA-transcription factor feedback motif, can give rise to sustained, tunable oscillations.

## Materials and methods

### Mathematical model

We model the interaction between a miRNA, miRNA sponge, mRNA, and transcription factor protein participating in a negative feedback loop. Our mathematical model is defined as follows, with parameter values in Table S1. We take the concentration of sponging RNA over time *t* as *C*(*t*), transcription factor mRNA as *F* (*t*), transcription factor protein as *P* (*t*), and miRNA as *M* (*t*). We denote basal rates of production of sponge RNA, miRNA, and transcription factor mRNA as *α*_*i*_ where *i ∈*{*C, M, F*}, respectively. We denote basal rates of degradation of sponge RNA, miRNA, transcription factor mRNA, and transcription factor protein as *δ*_*i*_ with *i ∈*{*C, M, F, P*}, respectively.

Inhibitory actions between two species *i* and *j* are assumed to follow mass-action kinetics (see [24] for a reference), with rate constant *k*_*ij*_ for (*i, j*) *∈* {(*C, M*), (*M, F*)} in the case of miRNA sponge repressing miRNA and miRNA repressing transcription factor mRNA, respectively.

We suppose that the rate of production of protein from mRNA for transcription follows a delayed linear relationship to the amount of mRNA, with an average translation rate of *k*_*P*_ per unit of mRNA. We represent time delays by *τ*_1_ and *τ*_2_ in this system to account for the transcription factor-mediated activation of transcription, and translation of mRNA into protein, respectively.

The interaction term between the transcription factor and its back-activation of miRNA production is defined by the following Hill function, as in similar models (e.g. [25]), such that:

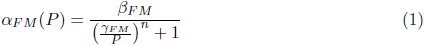

From a first-order mass-action kinetics formulation, we obtain the delay differential equations, with all derivatives taken with respect to time *t* signified by Ċ, Ṁ, Ḟ, Ṗ, as such:

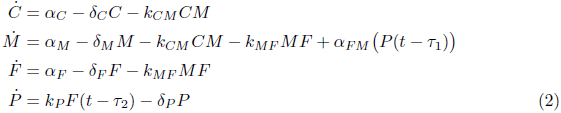

### Stochastic model

The reaction ‘events’ and the associated rates at which they occur in the stochastic version of our system are as described in System 2, with kinetic rate parameters on the right hand side, and a time delay indicated if present for that reaction. Each of the dynamic variables and parameters is as described above and in Table S1. The symbol Ø on the left side of a reaction indicates *de novo* synthesis, and on the right side of a reaction this indicates degradation.

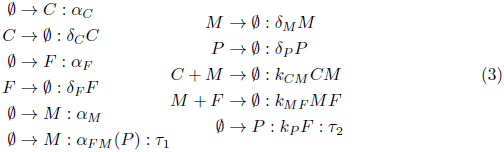

## Results

### Defining a miRNA-transcription factor feedback motif

The topology of the underlying network of interactions between RNA and proteins has a direct link to the system dynamics, and understanding this has led to wider predictions about the behaviour of biomolecules in the transcriptome [26–28]. For instance, extending these networks to include species of non-coding RNA, such as miRNA, which act to inhibit their predicted mRNA targets, has led to understanding of their functions in fine-tuning gene expression and maintaining bistable states[19, 29-31]. These transcriptome-wide studies have shown significant over-representation of specific miRNA-mRNA-protein subnetworks, representing distinct classes of feedback and feedforward motifs, each with unique intrinsic dynamical properties [32]. We consider an over-represented feedback motif involving a miRNA and transcription factor, as identified by Tsang et al. [32]. This motif is seen in an interaction between the E2F transcription factor and the miR-17/92 oncogenic cluster. Here, we will extend this by considering the effect of a miRNA sponge on network dynamics, depicted graphically in Fig 1 [33].

**Fig 1.**
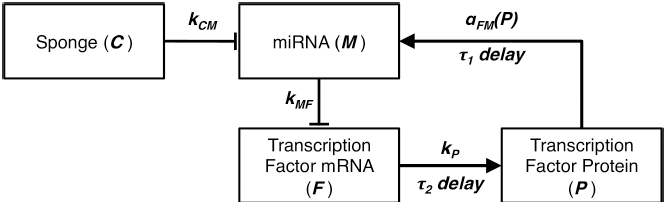
The miRNA sponge network considered. Directed arrows represent activation-type behaviour, and blunted arrows represent inhibitory behaviour. The system interconnections are overlayed with rate kinetic functions for each of the interactions and time delayed interactions are indicated by *τ*_1_ and *τ*_2_, yielding System 2.

We model this system mathematically by the set of equations outlined in the Materials and Methods. With this model, we analyse the long-term behaviour of this system via a stability analysis, and thereby study the properties of the unique equilibrium solution. The stability analysis gives an understanding of the qualitative behaviour of the system dynamics at equilibrium; for instance, whether it is stable steady state or oscillatory behaviour. A critical point at which the system changes from stable to oscillatory behaviour is known as a Hopf bifurcation, and the Hopf bifurcation theorem gives the necessary and sufficient conditions characterising whether these points occur, and the parameter values at these points. As per the derivation in Appendix S1, we apply the Hopf bifurcation theorem to show that for cases where the time delays are non-zero, there is a Hopf bifurcation when the sum of the time delays *τ*_1_ and *τ*_2_ exceeds some critical time *τ*_0_; resulting in a switch from asymptotic stability to an oscillatory steady state.

As a numerical illustration of this switch, consider the system for the following parameter values, chosen because they fall within a realistic range for known range parameters for mammalian cells as used in similar models (e.g. [34,35]):*α*_*C*_ = 1 mol *·* min^−1^, *δ*_*C*_ = 0.01 min^−1^, *α*_*F*_ = 1 mol *·* min^−1^, *δ*_*F*_ = 0.1 min^−1^, *α*_*M*_ = 1 mol *·* min^−1^, *δ*_*M*_ = 1 min^−1^, *k*_*P*_ = 10 mol *·* min^−1^, *δ*_*P*_ = 0.1 min^−1^, *k*_*CM*_ = 10 min^−1^ *·* mol^−1^, *k*_*MF*_ = 0.1 min^−1^ *·* mol^−1^, *β*_*F* *M*_ = 200 mol *·* min^−1^, *γ*_*F* *M*_ = 100 mol, and *n* = 8, with both cases of *τ*_1_ = *τ*_2_ = 0.5 min and *τ*_1_ = *τ*_2_ = 0.8 min as depicted in Fig 2A and B, respectively. These parameter values give a critical time *τ*_0_ of 1.43 min for which if *τ*_1_ + *τ*_2_ *> τ*_0_, there is an oscillatory solution, and when *τ*_1_ + *τ*_2_ *< τ*_0_ there is a steady state solution, as shown in Fig 2A and B.

**Fig 2.**
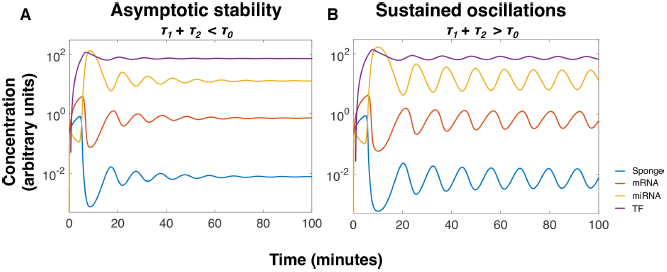
Increasing system delay past critical threshold induces steady oscillatory behaviour, traversing a Hopf bifurcation. Plots depict the effects of having *τ*_1_ + *τ*_2_ below (A) and above (B) the critical time threshold *τ*_0_ as derived above, based on the Hopf bifurcation theorem. Common parameter values used for this simulation are: *α*_*C*_ = 1 mol *·* min^−1^, *δ*_*C*_ = 0.01 min^−1^, *α*_*F*_ = 1 mol *·* min^−1^, *δ*_*F*_ = 0.1 min^−1^, *α*_*M*_ = 1 mol *·* min^−1^, *δ*_*M*_ = 1 min^−1^, *k*_*P*_ = 10 mol *·* min^−1^, *δ*_*P*_ = 0.1 min^−1^, *k*_*CM*_ = 10 min^−1^ *·* mol^−1^, *k*_*MF*_ = 0.1 min^−1^ *·* mol^−1^, *β*_*F* *M*_ = 200 mol *·* min^−1^, *γ*_*F* *M*_ = 100 mol, and *n* = 8, with *τ*_1_ and *τ*_2_ indicated as above.

### A novel mechanism for generating sustained oscillations

Our analysis shows that there is a critical sum of the two time delays, which is is a function of system parameter values, above which oscillatory behaviour emerges. This parametric dependence of the critical time may be exploited by biological systems to generate dynamic oscillatory behaviour, as although the parameters governing the kinetics and delays present in a biological system are largely fixed, rates of production and degradation vary significantly [35–38]. These may cause the system to move from an oscillatory state to a non-oscillatory state, or vice versa.

Transcriptional bursting is a phenomenon that has been observed across species for many genes, especially during development, whereby transcription rate is increased significantly in a ‘burst’ over a relatively short period of time [36]. As a descriptive example, we consider a time-varying value for *α*_*C*_, increasing it ten-fold from the baseline parameter values used in Fig 2. In this case, with a parameter value of *α*_*C*_ = 10 mol *·* min^−1^ the system has a critical time of *τ*_0_ = 0.62 min. The time delays *τ*_1_ and *τ*_2_ do not change with transcriptional bursting, and so the total delay remains*τ*_1_ + *τ*_2_ = 1 min, which is greater than the critical time during transcriptional bursting, so the system will exhibit oscillatory behaviour during bursting. To visualise this change, we show the system behaviour as *α*_*C*_ is increased ten-fold only transiently between simulation times 50 min and 150 min, and is 1 mol min^−1^ otherwise, in Fig 3. Here, oscillations are created dynamically and in a time-varying fashion, with their time to disappearance primarily determined by the miRNA sponge degradation rate.

**Fig 3.**
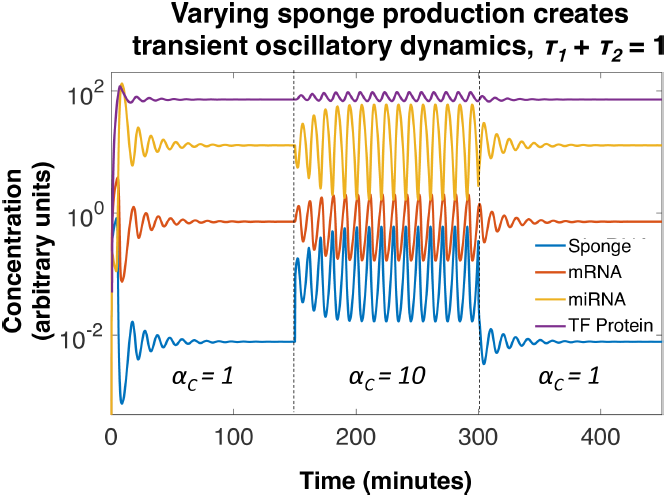
A time-varying *α*_*C*_ generates transient oscillatory behaviour. Here, a time varying value of *α*_*C*_ is used to illustrate the presence of a bifurcation. *α*_*C*_ is increased to 10 mol min^−1^ from an initial value of 1 mol min^−1^ between simulation time 150 min and 300 min, between which oscillatory behaviour is the absorbing state, and is reduced to 1 mol *·* min^−1^ otherwise, at which asymptotic stability predominates. Other parameter values are such that: *δ*_*C*_ = 0.01 min^−1^, *α*_*F*_ = 1 mol *·* min^−1^, *δ*_*F*_ = 0.1 min^−1^, *α*_*M*_ = 1 mol *·* min^−1^, *δ*_*M*_ = 1 min^−1^, *k*_*P*_ = 10 mol *·* min^−1^, *δ*_*P*_ = 0.1 min^−1^, *k*_*CM*_ = 10 min^−1^ *·* mol^−1^, *k*_*MF*_ = 0.1 min^−1^ *·* mol^−1^, *β*_*F* *M*_ = 200 mol *·* min^−1^, *γ*_*F* *M*_ = 100 mol, and *n* = 8, with *τ*_1_ = *τ*_2_ = 0.5 min as in Fig 2.

### Stochastic considerations

In the case where the number of molecules is small, as may occur in single cells with low copy numbers of these biomolecules, stochastic effects may predominate. In the stochastic setting, our system is no longer well described by the continuous variables written in System 2, but rather is better represented by a list of events that occur at discretised time steps, which we summarise in the Materials and Methods.

Moreover, because of the presence of non-zero time delays *τ*_1_ and *τ*_2_, this system exhibits non-Markovian behaviour, and therefore the stochastic behaviour may not follow the mean-field approximation by the ODE system in the long-term. That is, there may be oscillatory behaviour in the stochastic case for a parameter regime where the deterministic model does not predict oscillations [39]. This phenomenon, of *stochastic oscillations*, is one which we posit to be both significant and common among the behaviour of RNA networks, and has been thought to contribute to other oscillatory systems, such as the generation of circadian rhythms [40, 41], and in the Hes1 gene regulatory network [42].

To capture the potential for stochastic oscillations in our system, we simulate our system numerically using the dde23 Runge-Kutta based solver in Matlab, noting that conventional analytic approaches to this problem are intractable as they require deriving and solving the Langevin equations derived from the reactions in System 3. The algorithm we implement, described in Appendix S3 is based on the standard stochastic simulation algorithm by Gillespie, modified to handle the case of time-delayed reactions, also used for similar purposes such as delayed mRNA gene networks and chemical reaction networks [41,43, 44]. The original version of the Gillespie algorithm simulates reactions occurring instantaneously at randomly distributed times, weighted by their likelihood of occurring based on mass-action kinetics. In the modified algorithm, we account for time-delayed reactions by implementing a queuing system. If a time-delayed reaction is chosen to occur, it is not executed until a scheduled future time, determined by the length of time delay.

Fig 4 (left) depicts the results of a stochastic simulation for our system, showing oscillatory behaviour, with the overlayed mean field behaviour of *N* = 100 runs of the stochastic model (equivalent to the ODE solution). To study the periodicity of the stochastic signal, we take the discrete Fourier transform of the time dynamics, and analyse the power spectra for underlying modes. Shown in Fig 4 (right), this reveals a strong subcomponent of an underlying oscillatory mode for the stochastic simulations, whereas the deterministic model for this system with the same parameter values does not show this oscillatory mode.

**Fig 4.**
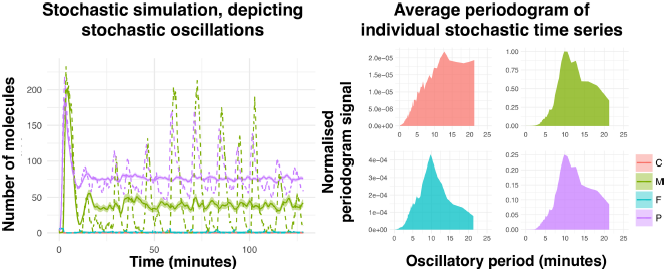
Stochastic system dynamics, showing an individual trace of mean behaviour and stochastic oscillations (left) and periodogram (right). Left: Averaged stochastic system dynamics do not show oscillations, but individual trajectories do. Dotted lines indicate an individual trajectory for a simulation, and bold lines are taken over an average of *N* = 100 runs, with standard error shaded around these lines. Right: Using the dynamics from stochastic simulations, we show the presence of underlying oscillatory modes, when the deterministic behaviour predicts asymptotic stability. Plots are of the average of *N* = 100 periodogram signal intensities, computed for each of the simulations of the stochastic model. Strong signal for an underlying oscillatory mode with period 10-15 minutes for the stochastic oscillations is evident, as corroborated by the individual series trace in Fig 4 (right). C refers to counts of miRNA sponge, M refers to counts of miRNA, F refers to counts of transcription factor mRNA, and P refers to counts of transcription factor protein. Parameter values used are the same as that of Fig 2, such that*α*_*C*_ = 1 mol *·* min^−1^, *δ*_*C*_ = 0.01 min^−1^, *α*_*F*_ = 1 mol *·* min^−1^, *δ*_*F*_ = 0.1 min^−1^, *α*_*M*_ = 1 mol *·* min^−1^, *δ*_*M*_ = 1 min^−1^, *k*_*P*_ = 10 mol *·* min^−1^, *δ*_*P*_ = 0.1 min^−1^, *k*_*CM*_ = 10 min^−1^ *·* mol^−1^, *k*_*MF*_ = 0.1 min^−1^ *·* mol^−1^, *β*_*F* *M*_ = 200 mol *·* min^−1^, *γ*_*F* *M*_ = 100 mol and *n* = 8, with *τ*_1_ = *τ*_2_ = 0.5 min, initial values chosen as five arbitrarily for all species.

## Discussion

Here, we have considered a common miRNA-transcription factor network motif extended to include a miRNA sponge. We have shown that in this system, without changing time delays or fixed kinetic parameters, oscillations can arise with simple changes in production and degradation rates of the miRNA sponge. In the stochastic case, oscillatory dynamics are more prevalent, and because of a change in bifurcation point location, are not seen in the deterministic limit for some cases. These results have implications that show how different types of non-coding RNA acting as miRNA sponges may generate dynamics not otherwise possible in a biological system.

### Different types of miRNA sponges confer different system behaviours

A key conclusion of this work is that different fixed kinetic properties of miRNA sponges will lead to different regimes and potentials for oscillatory dynamics in this network motif. As such, mapping these kinetic parameters to known information for the different species of RNA acting as miRNA sponges, we can hypothesise their effects, as depicted in Fig 5. For example, circRNA are differentiated from other species of ncRNA by their stability, as they do not have free ends, and so are not subject to the same RNA-se degrading enzymes [45, 46]. Based on our analysis, circRNA in this network motif acting as a miRNA sponge will push the steady state closer to oscillatory behaviour, potentially crucial to the maintenance of this state. In the same vein, recent work involving circRNA characterisation has shown, through knockdown experiments, that specific circRNA are heavily involved in neurogenesis, a process where such oscillatory behaviour is likely crucial [47, 48].

**Fig 5.**
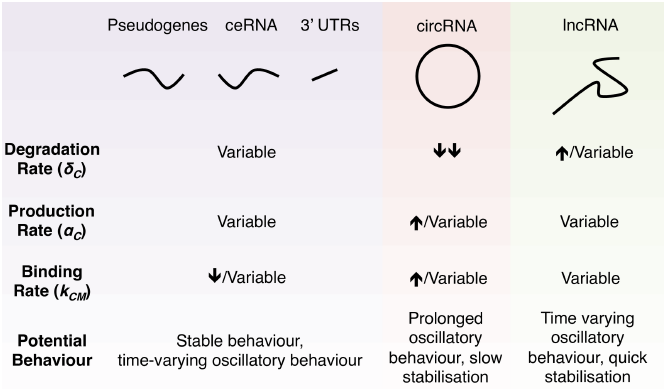
Summary of potential behaviours for different ncRNA acting as miRNA sponges in reaction network. Relationships between the dynamic parameters thought to occur for different ncRNA species functioning as miRNA sponges, and the effects of these parameter regimes has on system behaviour.

On the other hand, these results suggest that miRNA sponged by lncRNAs with a short half-life (as identified through a recent genome-wide analysis of lncRNA half-lives), are likely to exhibit greater stability and less propensity towards oscillatory behaviour [49]. In effect, these lncRNA, if produced in targeted bursts, may provide tight *temporal control* of oscillatory behaviour, perhaps crucial to regulating a switch between oscillatory and non-oscillatory behaviour, as in somitogenesis. It has also recently been shown that in mice, the lncRNA 116HG functions in the maintenance of circadian oscillations, as loss of this locus results in disruption of diurnal rhythms, although it is unknown if this is due to sponging functions [50].

### miRNA sponges in low copy number may be involved in the generation and maintenance of stochastic oscillations

As a result of the underlying biology of the interactions between the RNA, protein, and DNA species in our system, the mathematical description of this system involves delays.In practical terms, this results in the system exhibiting a phenomenon of stochastic oscillations, whereby stochastic system dynamics can exhibit oscillatory behaviour, even in a parameter regime where the deterministic solution does not.Translating this observation back into biologic terms, this result is particularly relevant for non-coding RNA, such as circRNA, which are thought to exist with low molecular counts, suggesting that oscillatory behaviour may be a more common feature of these RNA networks than would otherwise be predicted.

Next, we postulate the implications of extending the presented model to account for spatial differences in molecule concentration. Because the production of the biomolecules we consider is spatially organised within the cell, and they diffuse within the nucleus and cytoplasm, the concentration gradient of these molecules includes regions of very low numbers of molecules, up to much higher counts. As a result, the dynamics of any interacting network involving these species will differ along this gradient; encapsulating both stochastic and deterministic behaviours along different sub-regions in the cell. Regions closer to the edges of a diffusive boundary will have lower numbers of molecules, and, depending on parameter values, a greater propensity for stochastic oscillations. This may lead to a scenario in which there are steady state dynamics of the network at central regions of higher concentration, followed by disordered stochastic oscillatory behaviour as the biomolecules diffuse outward.Experiments, using techniques such as single RNA molecule tracking (e.g. [51, 52]) may be used to study how such spatial organisation may allow cells to generate oscillations at the behaviour of a cellular membrane; potentially facilitating motile behaviour, for example.

### Implications for ncRNA-based therapeutics

The results presented within this work also have implications for ncRNA-based therapeutic strategies. Fig S1 in Appendix S2 shows the key determinants of steady state levels for each of the species, through a parameter sensitivity analysis. Focussing on the values obtained for the system sensitivity to miRNA sponge parameters, we are able to infer the impact of a ncRNA therapeutic acting as a miRNA sponge on the network dynamics. For example, this shows that in order to decrease miRNA concentration, as opposed to increasing the binding kinetics of the sponge to the miRNA, we predict that it would be more effective to increase the production rate of the sponge (or introduce a higher concentration of miRNA sponge exogenously).

### A novel experimental paradigm

This work provides fertile ground for generating hypotheses regarding the functional roles of the various miRNA sponge species. However, we have done so within the confines of the limited evidence available at the present time. Characterisation of key kinetic parameters for miRNA sponge species, through the generation of synthetic forms, could provide ample substrate for more clearly elucidating their possible dynamics. Moreover, because we predict that these miRNA sponges may lead to oscillatory behaviour, the experimental design implemented must be robust enough to capture this. Instead of supposing *a priori* that there will be asymptotically stable dynamics, multiple time points with a sufficiently fine resolution must be considered to determine whether these oscillations are present. To optimise these time points for experimental assays, guidance should be sought from a theoretical model, with an analogous analysis as presented in this work.

## Conclusion

Overall, we have shown a novel paradigm by which oscillatory behaviour can emerge in RNA networks, via the actions of miRNA sponges. From this theoretical exploration, we have provided insight into the functional redundancy of miRNA sponges in different RNA configurations in nature. This, together with emerging knowledge of the roles of ncRNA, suggests that these species potentially have key implications for the behaviour of RNA networks in states of health and disease.

## Supporting information

**Appendix S1 Mathematical derivations.** Derivation of existence and uniqueness of system solution and stability analysis.

**Appendix S2 Parameter sensitivity analysis.** Analysis of system solution upon varying parameters through range of possible values.

**Appendix S3 Modified Gillespie Algorithm listing, pseudocode.** Pseudocode listing for the algorithm used for stochastic simulation, enabling the simulation of delayed reactions.

**Table S1 Parameter values.** Description of parameters and associated values as considered for sensitivity analysis.

**Fig S1 Partial rank correlation coefficients for each parameter value correlated with steady state values for modelled species**. Correlations are taken partial to all other parameter values. Parameter values were sampled using Latin hypercube sampling with 10^5^ points from the parameter space, and for each of these parameter combinations, steady state values were computed, from which the partial rank correlation coefficients can be presented. *C* refers to the steady state concentration of the miRNA sponge, *M* to the miRNA, *F* to the transcription factor mRNA, and *P* to the transcription factor protein.

**Fig S2 Partial rank correlation coefficients for each parameter value correlated with Euclidean norm of system steady state values.** Correlations are taken partial to all other parameter values. Parameter values were sampled using Latin hypercube sampling with 10^5^ points from the parameter space, and for each of these parameter combinations, steady state values were computed, from which the partial rank correlation coefficients can be presented.

**Fig S3 Logistic regression model coefficients for model parameters.** These are depicted with the associated 95% confidence interval for predicted coefficient. Model was trained as a classifier for whether regression would occur or not on 10^5^ parameter value combinations sampled from the space of possible values by Latin hypercube sampling. A positive coefficient indicates more likely to associate with existence of a bifurcation, and a negative coefficient indicates more likely to associate with global asymptotic stability.

**Fig S4 Partial rank correlation coefficient for each parameter value considered correlated with the critical time for which a bifurcation occurs.** Correlation is taken partial to all other parameters, for the cases in which a bifurcation is predicted to occur. Parameter values considered were selected using Latin hypercube sampling, using 10^5^ points in the parameter space, of which approximately 2300 had existence of a bifurcation.

## Acknowledgments

AD acknowledges funding support from Cancer Research UK.

## References

1. Glass L. Synchronization and rhythmic processes in physiology. Nature. 2001;410(6825):277–284.

2. Winfree AT. Biological rhythms and the behavior of populations of coupled oscillators. Journal of Theoretical Biology. 1967;16(1):15–42.

3. Mirollo RE, Strogatz SH. Synchronization of pulse-coupled biological oscillators. SIAM Journal on Applied Mathematics. 1990;50(6):1645–1662.

4. Wahl MB, Deng C, Lewandoski M, Pourquié O. FGF signaling acts upstream of the NOTCH and WNT signaling pathways to control segmentation clock oscillations in mouse somitogenesis. Development. 2007;134(22):4033–4041.

5. Serth K, Schuster-Gossler K, Cordes R, Gossler A. Transcriptional oscillation of lunatic fringe is essential for somitogenesis. Genes & Development. 2003;17(7):912–925.

6. Dale JK, Malapert P, Chal J, Vilhais-Neto G, Maroto M, Johnson T, et al. Oscillations of the snail genes in the presomitic mesoderm coordinate segmental patterning and morphogenesis in vertebrate somitogenesis. Developmental Cell. 2006;10(3):355–366.

7. Dequéant ML, Glynn E, Gaudenz K, Wahl M, Chen J, Mushegian A, et al. A complex oscillating network of signaling genes underlies the mouse segmentation clock. science. 2006;314(5805):1595–1598.

8. Pourquié O. The segmentation clock: converting embryonic time into spatial pattern. Science. 2003;301(5631):328–330.

9. Welsh DK, Logothetis DE, Meister M, Reppert SM. Individual neurons dissociated from rat suprachiasmatic nucleus express independently phased circadian firing rhythms. Neuron. 1995;14(4):697–706.

10. Goldbeter A. Computational approaches to cellular rhythms. Nature. 2002;420(6912):238–245.

11. Strogatz SH. From Kuramoto to Crawford: exploring the onset of synchronization in populations of coupled oscillators. Physica D: Nonlinear Phenomena. 2000;143(1-4):1–20.

12. Cooke J, Zeeman EC. A clock and wavefront model for control of the number of repeated structures during animal morphogenesis. Journal of theoretical biology. 1976;58(2):455–476.

13. Jiang YJ, Aerne BL, Smithers L, Haddon C, Ish-Horowicz D, Lewis J. Notch signalling and the synchronization of the somite segmentation clock. Nature. 2000;408(6811):475.

14. Baker RE, Schnell S, Maini P. A clock and wavefront mechanism for somite formation. Developmental Biology. 2006;293(1):116–126.

15. Herrgen L, Ares S, Morelli LG, Schröter C, Jölicher F, Oates AC. Intercellular coupling regulates the period of the segmentation clock. Current Biology. 2010;20(14):1244–1253.

16. Sturrock M, Hellander A, Matzavinos A, Chaplain MA. Spatial stochastic modelling of the Hes1 gene regulatory network: intrinsic noise can explain heterogeneity in embryonic stem cell differentiation. Journal of The Royal Society Interface. 2013;10(80):20120988.

17. Macnamara CK, Chaplain MA. Diffusion driven oscillations in gene regulatory networks. Journal of Theoretical Biology. 2016;407:51–70.

18. Phillips NE, Manning CS, Pettini T, Biga V, Marinopoulou E, Stanley P, et al. Stochasticity in the miR-9/Hes1 oscillatory network can account for clonal heterogeneity in the timing of differentiation. Elife. 2016;5.

19. Li JH, Liu S, Zhou H, Qu LH, Yang JH. starBase v2. 0: decoding miRNA-ceRNA, miRNA-ncRNA and protein–RNA interaction networks from large-scale CLIP-Seq data. Nucleic Acids Research. 2013;42(D1):D92–D97.

20. Thomson DW, Dinger ME. Endogenous microRNA sponges: evidence and controversy. Nature Reviews Genetics. 2016;17(5):272–283.

21. Paraskevopoulou MD, Georgakilas G, Kostoulas N, Reczko M, Maragkakis M, Dalamagas TM, et al. DIANA-LncBase: experimentally verified and computationally predicted microRNA targets on long non-coding RNAs. Nucleic Acids Research. 2012;41(D1):D239–D245.

22. Jeggari A, Marks DS, Larsson E. miRcode: a map of putative microRNA target sites in the long non-coding transcriptome. Bioinformatics. 2012;28(15):2062–2063.

23. Ebert MS, Sharp PA. MicroRNA sponges: progress and possibilities. RNA. 2010;16(11):2043–2050.

24. Horn F, Jackson R. General mass action kinetics. Archive for Rational Mechanics and Analysis. 1972;47(2):81–116.

25. Ingalls B, Mincheva M, Roussel MR. Parametric Sensitivity Analysis of Oscillatory Delay Systems with an Application to Gene Regulation. Bulletin of Mathematical Biology. 2017; p. 1–25.

26. Lee TI, Rinaldi NJ, Robert F, Odom DT, Bar-Joseph Z, Gerber GK, et al. Transcriptional regulatory networks in Saccharomyces cerevisiae. Science. 2002;298(5594):799–804.

27. Oates AC, Morelli LG, Ares S. Patterning embryos with oscillations: structure, function and dynamics of the vertebrate segmentation clock. Development. 2012;139(4):625–639.

28. Chaplain M, Ptashnyk M, Sturrock M. Hopf bifurcation in a gene regulatory network model: Molecular movement causes oscillations. Mathematical Models and Methods in Applied Sciences. 2015;25(06):1179–1215.

29. Volinia S, Galasso M, Costinean S, Tagliavini L, Gamberoni G, Drusco A, et al. Reprogramming of miRNA networks in cancer and leukemia. Genome Research. 2010;20(5):589–599.

30. Ryan BM, Robles AI, Harris CC. Genetic variation in microRNA networks: the implications for cancer research. Nature Reviews Cancer. 2010;10(6):389–402.

31. Lai X, Wolkenhauer O, Vera J. Understanding microRNA-mediated gene regulatory networks through mathematical modelling. Nucleic Acids Research. 2016;44(13):6019–6035.

32. Tsang J, Zhu J, van Oudenaarden A. MicroRNA-mediated feedback and feedforward loops are recurrent network motifs in mammals. Molecular Cell. 2007;26(5):753–767.

33. Aguda BD, Kim Y, Piper-Hunter MG, Friedman A, Marsh CB. MicroRNA regulation of a cancer network: consequences of the feedback loops involving miR-17-92, E2F, and Myc. Proceedings of the National Academy of Sciences. 2008;105(50):19678–19683.

34. Monk NA. Oscillatory expression of Hes1, p53, and NF-*κ*B driven by transcriptional time delays. Current Biology. 2003;13(16):1409–1413.

35. Schwanhäusser B, Busse D, Li N, Dittmar G, Schuchhardt J, Wolf J, et al. Global quantification of mammalian gene expression control. Nature. 2011;473(7347):337.

36. Suter DM, Molina N, Gatfield D, Schneider K, Schibler U, Naef F. Mammalian genes are transcribed with widely different bursting kinetics. Science. 2011;332(6028):472–474.

37. Cai L, Dalal CK, Elowitz MB. Frequency-modulated nuclear localization bursts coordinate gene regulation. Nature. 2008;455(7212):485.

38. Chen T, He HL, Church GM. Modeling gene expression with differential equations. In: Biocomputing’99. World Scientific; 1999. p. 29–40.

39. Liao S, Vejchodský T, Erban R. Tensor methods for parameter estimation and bifurcation analysis of stochastic reaction networks. Journal of The Royal Society Interface. 2015;12(108):20150233.

40. olde Scheper T, Klinkenberg D, Pennartz C, Van Pelt J. A mathematical model for the intracellular circadian rhythm generator. Journal of Neuroscience. 1999;19(1):40–47.

41. Bratsun D, Volfson D, Tsimring LS, Hasty J. Delay-induced stochastic oscillations in gene regulation. Proceedings of the National Academy of Sciences of the United States of America. 2005;102(41):14593–14598.

42. Galla T. Intrinsic fluctuations in stochastic delay systems: Theoretical description and application to a simple model of gene regulation. Physical Review E. 2009;80(2):021909.

43. Gillespie DT. Exact stochastic simulation of coupled chemical reactions. The Journal of Physical Chemistry. 1977;81(25):2340–2361.

44. Anderson DF. A modified next reaction method for simulating chemical systems with time dependent propensities and delays. The Journal of Chemical Physics. 2007;127(21):214107.

45. Enuka Y, Lauriola M, Feldman ME, Sas-Chen A, Ulitsky I, Yarden Y. Circular RNAs are long-lived and display only minimal early alterations in response to a growth factor. Nucleic Acids Research. 2016;44(3):1370–1383.

46. Gruner H, Cortés-López M, Cooper DA, Bauer M, Miura P. CircRNA accumulation in the aging mouse brain. Scientific Reports. 2016;6:38907.

47. Piwecka M, Glaˇzar P, Hernandez-Miranda LR, Memczak S, Wolf SA, Rybak-Wolf A, et al. Loss of a mammalian circular RNA locus causes miRNA deregulation and affects brain function. Science. 2017;357(6357):eaam8526.

48. Hanan M, Soreq H, Kadener S. CircRNAs in the brain. RNA Biology. 2017;14(8):1028–1034.

49. Clark MB, Johnston RL, Inostroza-Ponta M, Fox AH, Fortini E, Moscato P, et al. Genome-wide analysis of long noncoding RNA stability. Genome Research. 2012;22(5):885–898.

50. Powell WT, Coulson RL, Crary FK, Wong SS, Ach RA, Tsang P, et al. A Prader–Willi locus lncRNA cloud modulates diurnal genes and energy expenditure. Human molecular genetics. 2013;22(21):4318–4328.

51. Femino AM, Fay FS, Fogarty K, Singer RH. Visualization of single RNA transcripts in situ. Science. 1998;280(5363):585–590.

52. Park HY, Lim H, Yoon YJ, Follenzi A, Nwokafor C, Lopez-Jones M, et al. Visualization of dynamics of single endogenous mRNA labeled in live mouse. Science. 2014;343(6169):422–424.

